# Decoding COVID-19 mRNA Vaccine Immunometabolism in Central Nervous System: human brain normal glial and glioma cells by Raman imaging

**DOI:** 10.1101/2022.03.02.482639

**Authors:** H. Abramczyk, B. Brozek-Pluska, Karolina Beton

## Abstract

The paper presents the effect of COVID-19 mRNA (Pfizer/BioNT) vaccine on in vitro glial cells of the brain studied by means of Raman spectroscopy and imaging.. The results obtained for human brain normal and tumor glial cells of astrocytes, astrocytoma, glioblastoma incubated with the Covid-19 mRNA vaccine Pfizer/BioNT vaccine show alterations in the reduction-oxidation pathways associated with Cytochrome c.

We found that the Pfizer/BioNT vaccine down regulate the concentration of cytochrome c in mitochondria upon incubation with normal and tumorous glial cells. Concentration of oxidized form of cytochrome c in brain cells has been shown to decrease upon incubation the mRNA vaccine. Lower concentration of oxidized cytochrome c results in lower effectiveness of oxidative phosphorylation (respiration), reduced apoptosis and lessened ATP production. Alteration of Amide I concentration, which may reflect the decrease of mRNA adenine nucleotide translocator. Moreover, mRNA vaccine leads to alterations in biochemical composition of lipids that suggest the increasing role of signaling. mRNA vaccine produce statistically significant changes in cell nucleus due to histone alterations. The results obtained for mitochondria, lipid droplets, cytoplasm may suggest that COVID-19 mRNA (Pfizer/BioNT) vaccine reprograms immune responses. The observed alterations in biochemical profiles upon incubation with COVID-19 mRNA in the specific organelles of the glial cells are similar to those we observe for brain cancer vs grade of aggressiveness.

## 1. Introduction

The COVID-19 pandemics has witnessed an explosion in research in the field of immunometabolism that has revealed that similar mechanisms regulate the host response to infection, autoimmunity, and cancer. The new tools by Raman imaging we present in this paper raise exciting possibilities for new ways to understand pathways of our immune responses, recognize metabolites that regulates these pathways and suggest how we might use them to optimize vaccinations to stimulate the conditions of adaptive immune system.

The pandemic outbreak in 2019 by the SARS-COV-2 virus generating acute respiratory syndrome caused 230 418 451 confirmed cases of COVID-19, including 4 724 876 deaths, reported to WHO. As of 23 September 2021, a total of 5 874 934 542 vaccine doses have been administered [1]. In response to the urgent need for a vaccine, pharmaceutical companies including Pfizer/BioNT in 2020 proposed vaccines based on mRNA technology. The Pfizer/BioNT vaccine (BNT162b2) is more than 90% effective against COVID-19 [2].

In order to enter the host cells, the SARS-COV-2 virus uses the S surface protein, the so-called spike protein (spike S protein). Vaccines based on mRNA technology are designed to produce antibodies to the spike protein. mRNA vaccines are vaccines in which ribonucleic acid (RNA) is used as a template for the production of viral proteins. These proteins are designed to trigger the production of antibodies, which are then transferred to the host’s immune system.

In the paper, we present effect of mRNA vaccines on the glial brain cells that are involved in tumor microenvironment infiltration by using a novel non-invasive tool such as Raman imaging. Here we demonstrate that Raman imaging reveal new expanses on the role of basic mechanisms of cancer pathology and effect of mRNA vaccines. This approach can monitor interactions in tumor microenvironment and mechanisms related to immune response.

Raman spectroscopy and imaging enabling quantitative and non-invasive monitoring of intracellular changes without the need of using external markers. Traditional methods of molecular biology require the destruction of cell membranes and the isolation of intracellular components to study the biochemical changes inside cells. In Raman imaging, we do not need to destroy cells to learn about biochemical composition of intracellular structures (cell organelles). Tracking alterations in biochemical composition in separate organelles is extremely valuable in establishing molecular mechanism of cancer development and mechanisms of infections. Until now, no technology has proven effective for detecting concentration of specific compounds in separate cell organelles. Therefore, existing analytical technologies cannot detect the full extent of biolocalization inside and outside specific organelles.

We will concentrate on normal and tumor glial cells upon incubation with mRNA vaccine. The reason is that cancer diseases are the most serious cause of death, exceeding heart disease, strokes, pneumonia and COVID-19. Although at the moment there is no vaccine against most cancers, rapid development of mRNA vaccines may help in development anticancer vaccines.

The announcement of effective and safe vaccines for COVID-19 has been greeted with enthusiasm. The vaccines currently used in the global vaccination campaign (3.36 billion doses have been administered across 180 countries, according to data collected [https://www.bloomberg.com/graphics/covid-vaccine-tracker-global-distribution/].

While COVID-19 vaccines bring potential hope for a return to some kind of normality, many of the fundamental mechanisms by which mRNA vaccines induce strong responses are still incompletely understood and should be continued [2]. The mRNA vaccine encoding the COVID-19 spike (S) protein encapsulated in lipid nanoparticles gain entry into dendritic cells (DCs) at the injection site or within lymph nodes, resulting in production of high levels of S protein.

Nevertheless, there is still much to learn. It is not clear which cell specific activation contributes the most to vaccine efficacy and what activation may inhibit the generation of adaptive immunity or lead to poor tolerability of the vaccine [3].

There are controversies on harmful effects from spike S protein produced by COVID-19 vaccination and long-term effects. Researchers have warned that Pfizer-BioNTech’s coronavirus disease 2019 (COVID-19) vaccine induces complex reprogramming of innate immune responses that should be considered in the development and use of mRNA-based vaccines. There are also controversies on biodistribution of mRNA vaccines. It has been reported [4] that intramuscular vaccines (which Pfizer/BioNT vaccine is) in macaques (a type of monkey) remain near the site of injection (the arm muscle) and local lymph nodes, where white blood cells and antibodies are produced to protect from disease. The lymphatic system, lymph nodes also clean up fluids and remove waste materials. Similar results were obtained for a mRNA vaccine against H10N8 and H7N9 influenza viruses in mice [4]. However, recent results on interactions between the immune system and the viral proteins that induce immunity against COVID-19 may be more complex than previously thought [5]. Evidence has been found that spike S protein of COVID-19 has remain not only near the site of injection, but also circulate in the blood. COVID-19 proteins were measured in longitudinal plasma samples collected from 13 participants who received two doses of mRNA-1273 vaccine. 11 of 13 participants showed detectable levels of COVID-19 protein as early as day one upon first vaccine injection. Clearance of detectable COVID-19 protein correlated with production of IgG and IgA [6].

In the view of the recent results it is important to be aware that the spike S protein produced by the new COVID-19 mRNA vaccines may also directly affect the host cells with the long-term consequences. Thus, one should monitor biodistribution and location of spike S protein from mRNA vaccines and the effects of the COVID-19 spike S protein on human host cells in vitro and appropriate experimental animal models.

In this paper we will concentrate on central nervous system (CNS) because in addition to pneumonia and acute respiratory distress, COVID-19 is associated with a host of symptoms that are related to the CNS, including loss of taste and smell, headaches, twitching, seizures, confusion, vision impairment, nerve pain, dizziness, impaired consciousness, nausea and vomiting, hemiplegia, ataxia, stroke and cerebral hemorrhage [7].

It is unclear whether severe acute respiratory syndrome coronavirus, which causes COVID-19, can enter the brain. It has been postulated that some of the symptoms of COVID-19 may be due to direct actions of the virus on the CNS; for example, respiratory symptoms could be in part due to COVID-19 invading the respiratory centers of the brain [8,9]. Encephalitis has also been reported in COVID-19, and could be a result of virus or viral proteins having entered the brain [7,10].

COVID-19 mRNA has been recovered from the cerebrospinal fluid [11], suggesting it can cross the blood–brain barrier (BBB). Other coronaviruses, including the closely related SARS virus that caused the 2003–2004 outbreak, are able to cross the BBB [12], and COVID-19 can infect neurons in a Brain Sphere model [13]. However, COVID-19 could induce changes in the CNS without directly crossing the BBB, as COVID-19 is associated with a cytokine storm, and many cytokines cross the BBB to affect CNS function [9]. It has been found that COVID-19 reaches the brain, infects astrocytes and triggers neuropathological changes that contribute to the structural and functional alterations in the brain of COVID-19 patients [14]. The researchers raised a concern that the lipid nanoparticles (LNPs) that can diffuse quickly, could potentially gain access to the central nervous system (CNS) through the olfactory bulb or blood. However, this needs to be determined with further study. Also, the role of innate memory responses to LNPs needs to be studied [15].

Visualization of alterations in single cells upon delivery of mRNA vaccines would help evaluate the efficacy of candidate formulations and aid their rational design for preclinical and translational studies. Here, we show that Raman imaging allows for quantitative, and non-invasive monitoring response to mRNA vaccine in specific organelles without any labeling.

In this paper we will study implications for possible consequences of COVID-19 mRNA vaccine (Pfizer/BioNT BNT162b2) on the central nervous system (CNS). We have already studied in detail the biochemical alterations in specific organelles of brain cells vs cancer aggressiveness.[16,17] Thus, we can compare the effect of mRNA on normal and cancer cells with the effect of cancer aggressiveness. As far as we know the mRNA Pfizer vaccine has not been tested for patients suffering of cancer. Therefore this contribution will help monitoring responses in host brain cells similar to a viral infection, because the incubation with COVID-19 mRNA vaccine mimics COVID-19 infection, but instead of the whole virus, only one key protein S for the immune response is synthesized, without causing COVID-19 infection.

We will study human brain normal glial cells and glioma cells in vitro: normal human astrocytes (Clonetics NHA), human astrocytoma CCF-STTG1 (ATTC CRL-1718) representing mildly aggressive brain tumor and human glioblastoma cell line U87-MG (ATCC HTB-14) representing highly aggressive brain tumor by Raman imaging. mRNA vaccine mimics COVID-19infection, but instead of the whole virus, only one key protein S for the immune response is synthesized, without causing COVID-19 infection.

We will monitor the effect of the mRNA vaccine on biodistribution of different chemical components, particularly cytochrome c, in the specific organelles of a cell: nucleus, mitochondria, lipid droplets, cytoplasm and membrane.

In the presented study we will identify dynamics and biochemical composition of the organelles through characteristic Raman spectra upon injection of mRNA vaccine and incubation with the vaccine in vitro cells.

We will show also that Raman spectroscopy and Raman imaging are competitive clinical diagnostics tools for cancer diseases linked to mitochondrial dysfunction and are a prerequisite for successful pharmacotherapy of cancer.

In this paper we explore alterations in reduction-oxidation pathways related to Cyt c in human brain normal and tumor cells upon incubation in vitro with COVID-19 vaccine (Pfizer/ BioNT BNT162b2).

## 2. Materials and Methods

### 2.1. Reagents

Cytochrome c (C 2506) was purchased from Sigma Aldrich (Poland).

### 2.2 In vitro cells culturing and incubation with vaccine

The studies were performed on normal human astrocytes (Clonetics NHA), human astrocytoma CCF-STTG1 (ATTC CRL-1718) and human glioblastoma cell line U87-MG (ATCC HTB-14) purchased from Lonza (Lonza Walkersville. Inc., Walkersville, MA, USA) and American Type Culture Collection (ATCC), respectively. The NHA cells were maintained in Astrocyte Medium Bulletkit Clonetics (AGM BulletKit, Lonza CC-3186) and Reagent Pack (Lonza CC-5034) without antibiotics in a humidified incubator at 37 °C and 5% CO_2_ atmosphere. The U87MG cells were maintained in Eagle’s Minimal Essential Medium with L-glutamine (ATCC 30-2003) supplemented with 10% fetal bovine serum (ATCC 30-2020) without antibiotics in a humidified incubator at 37 ° C and 5% CO_2_ atmosphere. The CRL-1718 cells were maintained in RPMI 1640 Medium (ATCC 30-2001) supplemented with 10% fetal bovine serum (ATCC 30-2020) without antibiotics in a humidified incubator at 37 °C and 5% CO_2_ atmosphere. Cells were seeded on CaF_2_ window (Crystran Ltd., Poole, UK; CaF_2_ Raman grade optically polished window 25 mm diameter × 1 mm thick, no.CAFP25-1R, Poole, UK) in a 35 mm Petri dish at a density of 5 × 104 cells per Petri dish. Upon 24 h of incubation of cells in pure culture medium, cells were supplemented with the vaccine at various time and concentration variants. The COVID-19 mRNA (Pfizer/BioNT vaccine was diluted in 1,8 mL of 0,9% sodium chloride. The total volume of used Petri dishes was 3 mL, so that we added 180 µL of the diluted vaccine to 3 mL of pure culture medium The real dose of the vaccine that is administered to patients is equal to 2.25 mL/6= 0.3 mL corresponding to 30 µg per dose [18]. It is difficult to estimate the volume of the body in which the vaccine is diluted in real patients. The doses of 1 µL/mL and 60 µL/mL we used correspond to the dose given during the vaccination of real patients assuming the local distribution in the place of injection and around 100 times higher when we use the volume of the fluids in the human body). Before Raman examination, cells were fixed with 4% formalin solution (neutrally buffered) for 10 minutes and kept in phosphate-buffered saline (PBS, no. 10010023, Gibco) during the experiment.

### 2.3. Raman imaging and spectroscopy

Raman spectroscopy is an analytical technique where inelastic scattered light is used to obtain the information about the vibrational energy of analyzed samples. In the vast majority of scattering events, the energy of the molecule is unchanged after its interaction with the photon; and therefore the wavelength, of the scattered photon is equal to that of the incident photon. This is called elastic (energy of scattering particle is preserved) or Rayleigh scattering and is the dominant process during interaction of photon with the molecule. In a much rarer event (approximately 1 in 10 million photons) Raman scattering occurs, which is an inelastic scattering process with a transfer of energy between the molecule and scattered photon. If the molecule gains energy from the photon during the scattering (excited to a higher vibrational level) then the scattered photon loses energy and its wavelength increases which is called Stokes Raman scattering. Inversely, if the molecule loses energy by relaxing to a lower vibrational level the scattered photon gains the corresponding energy and its wavelength decreases; which is called Anti-Stokes Raman scattering. Quantum-mechanically, Stokes and Anti-Stokes are equally probable processes. However, with an ensemble of molecules, the majority of molecules will be in the ground vibrational level (Boltzmann distribution) and Stokes scattering is statistically more probable process. In consequence, the Stokes Raman scattering is always more intense than the Anti-Stokes component and for this reason, it is nearly always the Stokes Raman scattering that is measured wherewithal Raman spectroscopy. Scheme 1 presents illustration of Rayleigh, Stokes and Anti-Stokes Raman Scattering.

**Scheme 1.**
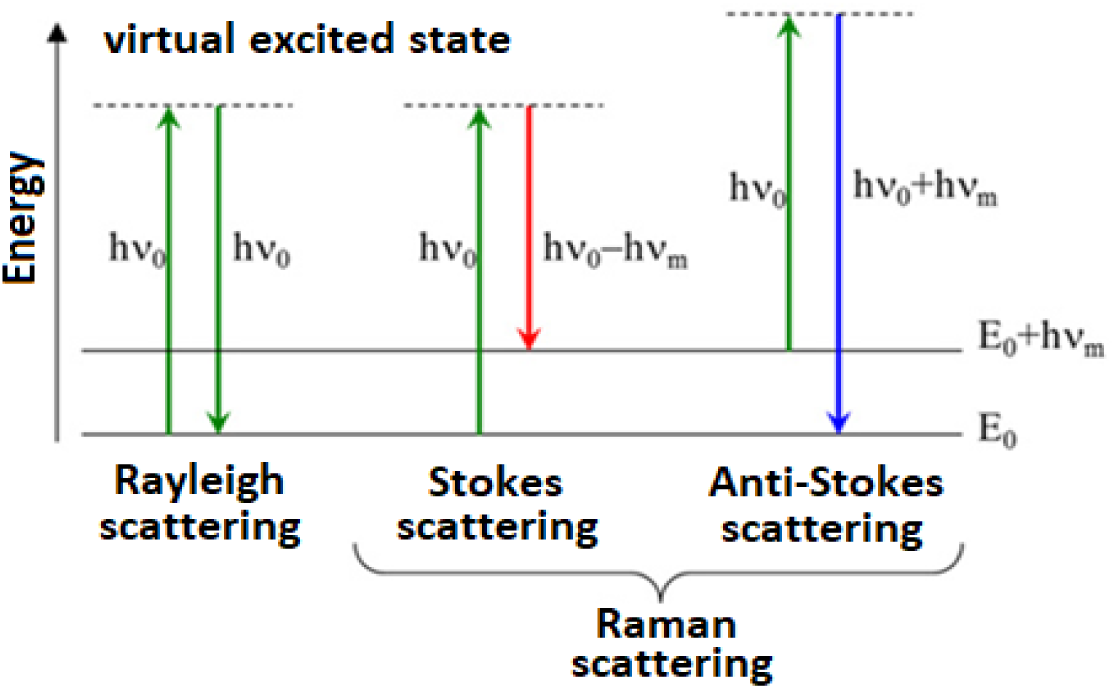
Schematic presentation of scattering phenomena.

Raman imaging is a technique based on Raman scattering allowing not only a single spectrum acquisition characteristic for a single point of the sample but also the analysis of vibrational spectra of any sample area. The imaging mode allows the analysis of distribution of different chemical molecules inside the sample. Using algorithms such as Cluster Analysis (see section 2.4) based on 2D data obtained by using Raman imaging make possible to create Raman maps to visualize cell’s substructures: nucleus, mitochondria, lipid structures, cytoplasm, cell membrane and learn about their biocomposition.

**Scheme 2.**
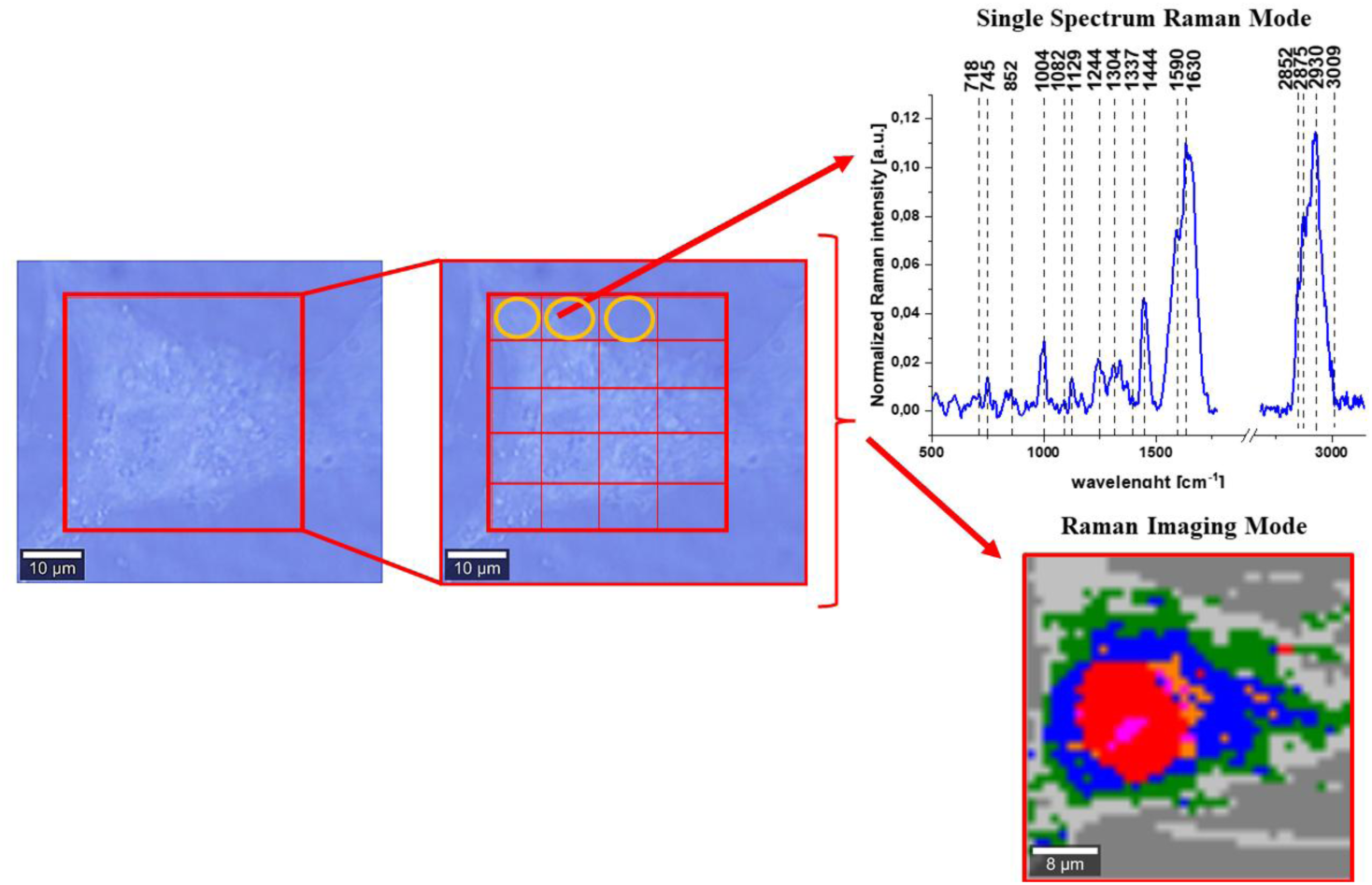
Schematic comparison of Raman single spectra and Raman imaging modes of data acquisition.

Raman spectra and images were recorded using a confocal Raman microscope (WITec (alpha 300 RSA+), Ulm, Germany) in the Laboratory of Laser Molecular Spectroscopy, Lodz University of Technology, Poland. The Raman microscope consisted of an Olympus microscope (Olympus Düsseldorf, Germany) a UHTS (Ultra-High-Throughput Screening) monochromator (WITec, Ulm, Germany) and a thermoelectrically cooled CCD camera ANDOR Newton DU970N-UVB-353 (EMCCD (Electron Multiplying Charge Coupled Device, Andor Technology, Belfast, Northern Ireland) chip with 1600 × 200 pixel format, 16 µm dimension each) at −60° C with full vertical binning. A 40× water immersion objective (Zeiss, W Plan-Apochromat 40×/1.0 DIC M27 (FWD = 2.5 mm), VIS-IR) was used for cell lines measurements. The excitation laser at 532 nm was focused on the sample to the laser spot of 1 µm and was coupled to the microscope via an optical fiber with a diameter of 50 µm. The average laser excitation power was 10 mW, and the collection time was 0.5 and 1 s for Raman images. Raman images were recorded with a spatial resolution of 1 × 1 µm. The Raman microspectrometer was calibrated every day prior to the measurements using a silica plate with a maximum peak at 520.7 cm^-1^.

### 2.4. Data processing

Data acquisition and processing were performed using WITec Project Plus software. The background subtraction and the normalization (model: divided by norm (divide the spectrum by the dataset norm)) were performed by using Origin software. The normalization model: divided by norm was performed according to the formula:

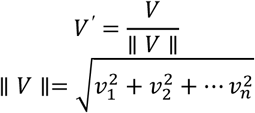

where:

*v*_*n*_ is the n^th^ V values.

The normalization was performed for low (500-1800 cm^-1^) and high (2600-3500 cm^-1^) frequency spectral regions separately.

For each organelle of a cell we have recorded hundreds of Raman spectra (because we have recorded one single Raman spectrum for each point of the imaging area marked in red in the microscopic image of the Scheme 2 with the resolution of 1 µm. We used Cluster Analysis method, implemented in WITec project software to calculate the average Raman spectra.

### 2.5. Cluster analysis

Spectroscopic data were analyzed using Cluster Analysis method. Briefly Cluster Analysis is a form of exploratory data analysis in which observations are divided into different groups that have some common characteristics – vibrational features in our case. Cluster Analysis constructs groups (or classes or clusters) based on the principle that: within a group the observations must be as similar as possible, while observations belonging to different groups must be different.

The partition of n observations (x) into k (k≤n) clusters S should be done to minimize the variance (Var) according to the formula:

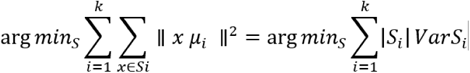

where

*μ*_*i*_ is the mean of points *S*_*i*_.

Raman maps presented in the manuscript were constructed based on principles of Cluster Analysis described above. Number of clusters was 7 (the minimum number of clusters characterized by different average Raman spectra, which describe the variety of the inhomogeneous biological sample).

## 3. Results

To learn about alterations in biochemical composition in cell organelles by methods of conventional molecular biology one have to disrupt a cell to release the cellular structure to estimate fractions that are enriched with specific organelles. Using Raman imaging we do not need to break cells to learn about the localization, distribution and bio-chemical composition of specific compounds in different organelles. To properly address alterations in single brain cells upon incubation with COVID-19 Pfizer/BioNT vaccine, we systematically investigated how Raman spectroscopy and Raman imaging monitor responses to the vaccine in specific organelles.

Figure 1 shows the Raman image of a single cell of glioblastoma (U-87 MG) of highly aggressive brain tumor incubated with mRNA-based Pfizer/BioNT vaccine (dose 60 µL/mL) for 96 hours and corresponding Raman imagines of specific organelles. The Raman images were created by K-means cluster analysis using 7 clusters. The blue color represents lipids including rough endoplasmic reticulum and lipid droplets (filled with retinoids), the orange color represents lipid droplets (filled with triacylglycerols of monounsaturated type (TAG), magenta color represents mitochondria, red color represents nucleus, green-cytoplasm, and light grey – membrane (the dark grey color corresponds to cell environment).

**Figure 1.**
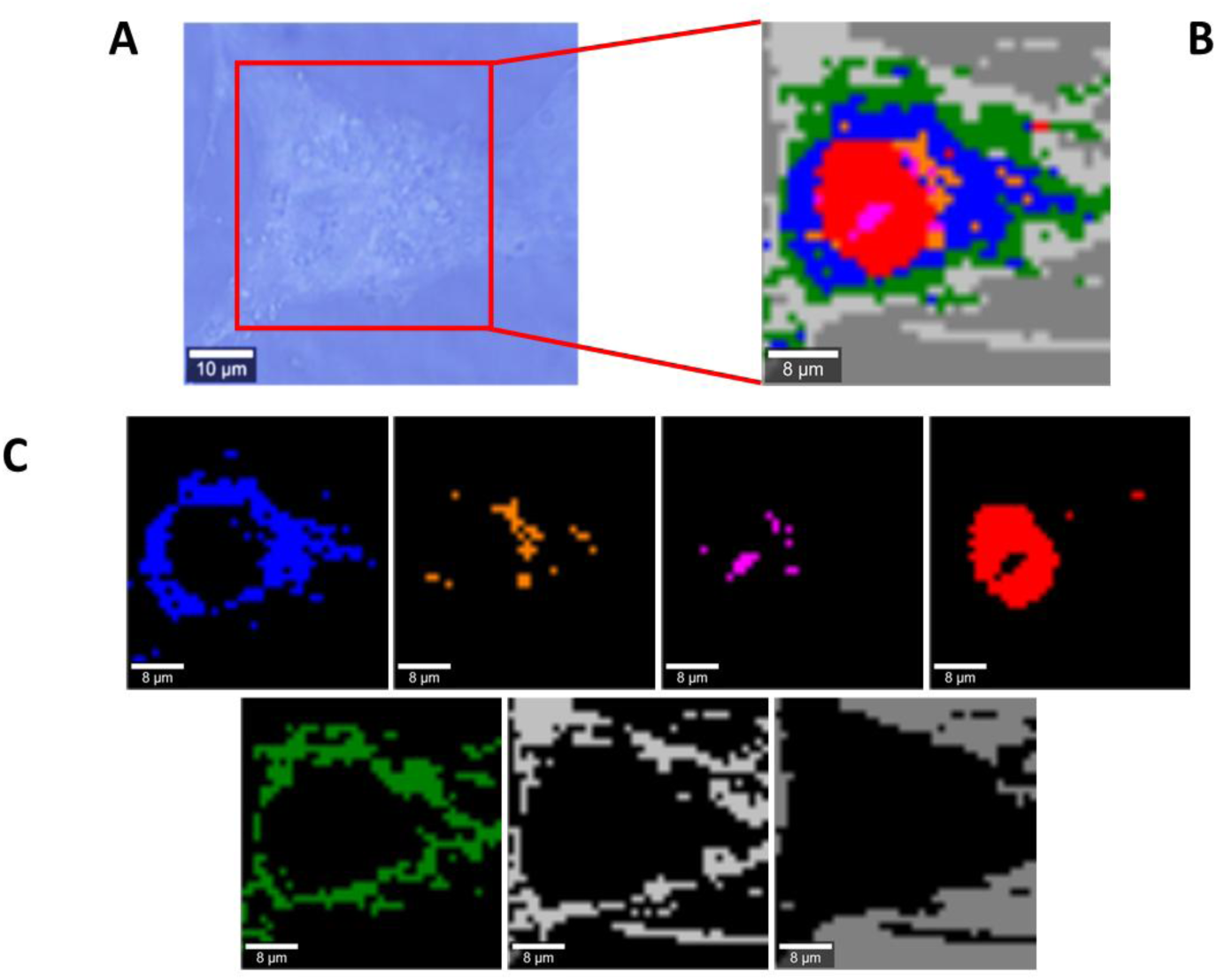
Microscopy image (A), Raman image of glioblastoma (U-87 MG) cell (30×20 µm, resolution 1.0 µm) incubated with Pfizer/BioNT vaccine (dose 60 µL/mL) for 96 hours (B) and Raman imagines of specific organelles: lipids and lipid droplets (blue and orange), mitochondria (magenta),nucleus (red), cytoplasm (green), membrane (light grey), cell environment (dark grey) at 532 nm.

Figure. 2 shows the Raman image of a single cell of astrocytoma (CRL-1718) of mildly aggressive brain tumour and corresponding Raman images of specific organelles incubated with mRNA-vaccine from Pfizer/BioNT (dose 60 µL/mL) for 96 hours. The Raman images were created by K-means cluster analysis using 7 clusters with the same coding colours as in Fig.1.

**Figure 2.**
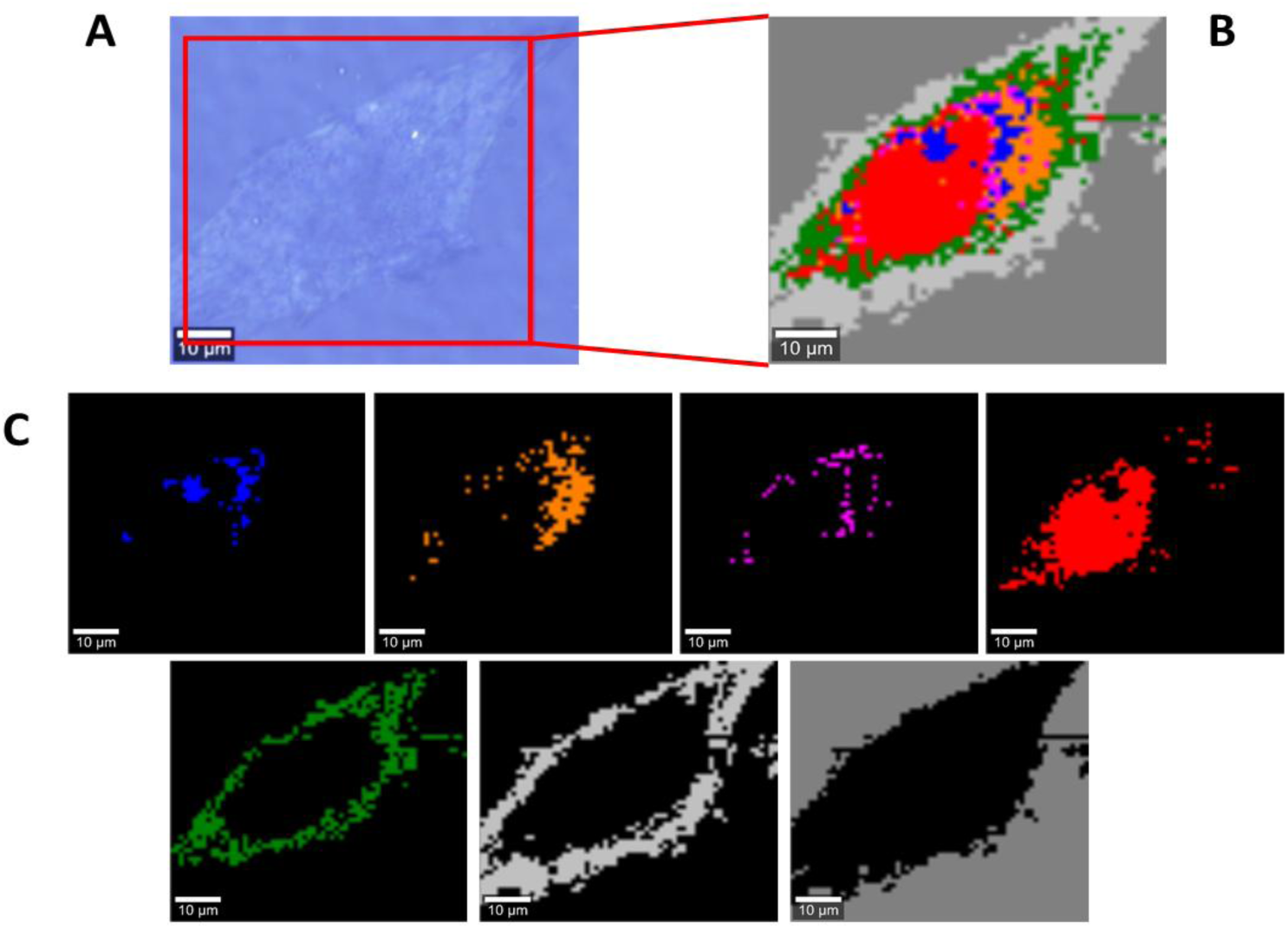
Microscopy image (A), Raman image of astrocytoma (CRL1718) cell (65×60 µm, resolution 1.0 µm) incubated with Pfizer/BioNT vaccine (dose 60 µL/mL) for 96 hours (B) and Raman imagines of specific organelles: lipids and lipid droplets (blue and orange), mitochondria (magenta),nucleus (red), cytoplasm (green), membrane (light grey), cell environment (dark grey) at 532 nm.

Figure. 3 shows the Raman image of a single normal cell of astrocyte (NHA) and corresponding Raman images of specific organelles. The Raman images were created by K-means cluster analysis using 7 clusters with same the coding colours as in Fig. 1.

**Figure 3.**
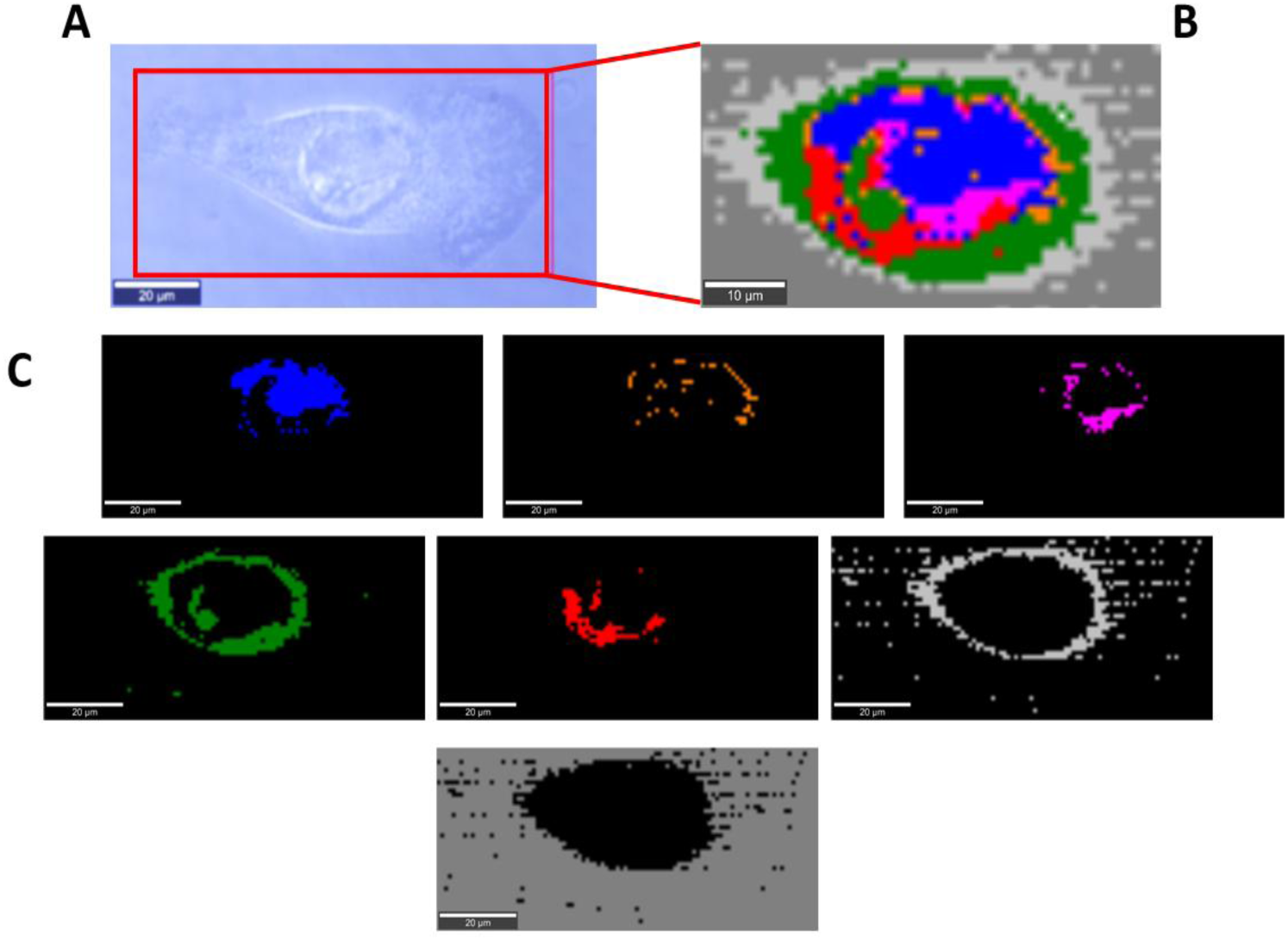
Microscopy image (A), Raman image of astrocyte (NHA) cell (100×45 µm, resolution 1.0 µm) incubated with Pfizer/BioNT vaccine (dose 60 µL/mL) for 96 hours (B) and Raman imagines of specific organelles: lipids and lipid droplets (blue and orange), mitochondria (magenta),nucleus (red), cytoplasm (green), membrane (light grey), cell environment (dark grey) at 532 nm.

To properly address alterations in brain tumor cells upon incubation with COVID-19 vaccine (Pfizer/BioNT) we systematically investigated how the Raman imaging and spectroscopy monitor response of in vitro human brain normal cells and cells of different aggressiveness.

### 3.1. Mitochondria-mRNA

Figure. 4 shows the effect of the Pfizer/BioNT vaccine on mitochondria in glioblastoma cells (U87 MG), astrocytoma (CRL-1718) and normal cells of astrocyte (NHA) by Raman imaging. Fig 4 shows a comparison of the average Raman spectra (normalized by vector norm) for mitochondria at 532 nm excitation with and without mRNA vaccine.

**Figure 4.**
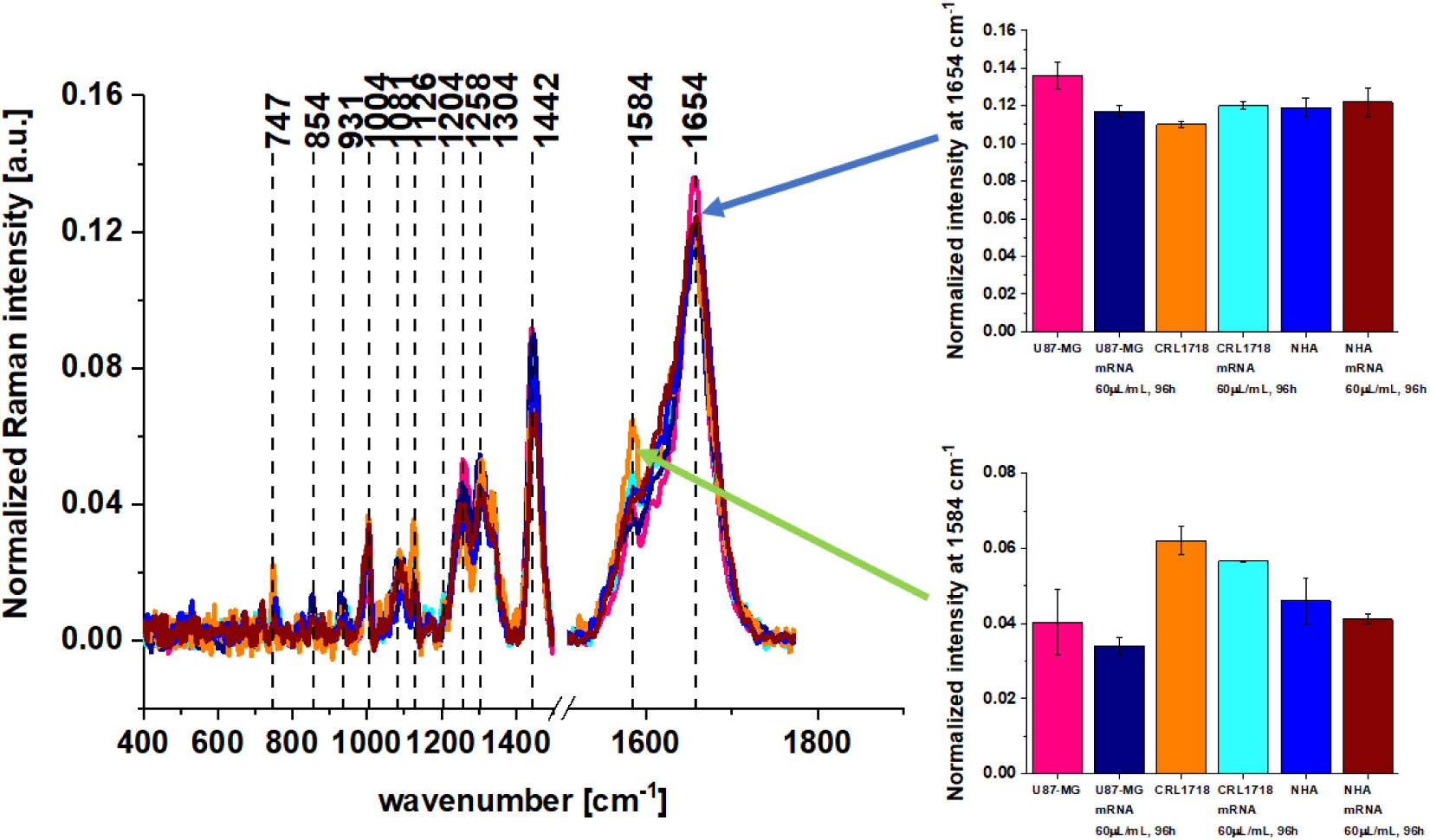
The effect of the Pfizer/BioNT vaccine from on mitochondria in glioblastoma cells (U87 MG)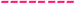, glioblastoma cells (U87 MG) upon incubation with Pfizer/BioNT vaccine for 96h 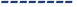, astrocytoma (CRL-1718) 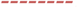, astrocytoma cells (CRL1718) upon incubation with Pfizer/BioNT vaccine for 96h 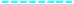, normal astrocytes (NHA) 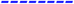, and normal astrocytes (NHA) upon incubation with Pfizer/BioNT vaccine for 96h 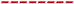 (number of cells for each cell’s type: 3, number of Raman spectra for each single cell: minimum 1600, excitation wavelength: 532 nm, laser power: 10 mW, integration time: 1.0sec).

Detailed inspection into Fig. 4 demonstrates that the most significant changes occur at 1584 cm^-1^. The peak at 1584 cm^-1^ represents the “redox state Raman marker” of Cyt c concentration. Recently we demonstrated that this Raman vibration can serve as a sensitive indicators cytochrome c concentration and correlates with cancer aggressiveness [16]. We showed that the Raman intensity of the band at 1584 cm^-1^ corresponding to the concentration of cytochrome c in mitochondria of a cell in vitro decreases with brain tumor aggressiveness.[16,17]

Briefly, cytochromes are classified on the basis of their lowest electronic energy absorption band in their reduced state. Therefore, we can distinguish cytochrome P450 (450 nm), cytochrome c (550 nm), cytochromes b (≈565 nm), cytochromes a (605 nm). The cytochromes are localized in the electron transport chain in the complex known as complex III or Coenzyme Q – Cyt C reductase, sometimes called also the cytochrome bc1 complex (cytochrome b, cytochrome c1). Cytochrome c, which is reduced to cytochrome c Fe^2+^ by the electron from the complex III to complex IV, where it passes an electron to the copper binuclear center, being oxidized back to cytochrome c (cyt c Fe^+3^). Complex IV is the final enzyme of the electron transport system. The complex IV contains two cytochromes a and a3 and two copper centers.

Until now, no technology has proven effective for detecting Cyt c concentration in specific cell organelles. Existing analytical technologies such as enzyme-linked immunosorbent assays (ELISA), Western blot, high performance liquid chromatography (HPLC), spectrophotometry and flow cytometry cannot detect the full extent of Cyt c localization inside and outside specific organelles. Therefore, none of the methods used to control Cyt c concentration can provide direct evidence about the role of cytochrome c in apoptosis and oxidative phosphorylation, because they are not able to monitor the amount of cytochrome in specific organelles such as mitochondria, cytoplasm, or extracellular matrix. Raman imaging does not need to disrupt cells to open yjrm to release the cellular structures to learn about their biochemical composition. Recently we showed [16,19] that the concentration of Cyt c in cancer tissues increases with cancer aggressiveness. It indicates that the netto concentration of Cyt c in mitochondria is higher than release to cytoplasm. This finding reflects the dual face of Cyt in life and death decisions: apoptosis and oxidative phosphorylation. The balance between cancer cells proliferation (oxidative phosphorylation) and death (apoptosis) decide about level of cancer development. The Cyt c concentration in mitochondria as a function of cancer aggressiveness reflects its contribution to oxidative phosphorylation and apoptosis.[16,19] In contrast, in single cells in vitro where the interactions with the extracellular matrix are eliminated the trend is opposite than in the cancer tissues. The biochemical results obtained by Raman imaging showed that human single cells in vitro demonstrate a redox imbalance by downregulation of cytochrome *c* in cancers.

One can see from Figure 4 that the Raman signal of oxidized cytochrome c is the strongest for astrocytoma control cells and the weakest one for high-grade glioblastoma (U-87 MG). It indicates that the processes of oxidative phosphorylation and apoptosis decrease with cancer aggressiveness.

Fig. 4 also shows significant changes at 1654 cm^-1^ corresponding to Amide I vibrations. The intensity of the band at 1654 cm^-1^ decreases for glioblastoma U87 MG upon incubation with mRNA. It has been reported that changes in mitochondrial membrane potential favour functional deterioration of the adenine nucleotide translocator (ANT), which belongs to mitochondrial carrier family [20]. As ANT represents about 10 % of proteins in mitochondria the observed in Fig. 4 decrease upon incubation with mRNA may reflect this deterioration. Figure. 5 shows a comparison of the average Raman spectra (normalized by vector norm) obtained for mitochondria in glioblastoma cells (U87 MG), astrocytoma (CRL-1718) and normal cells of astrocyte (NHA) with and without mRNA vaccine for 532 nm excitation.

**Figure 5.**
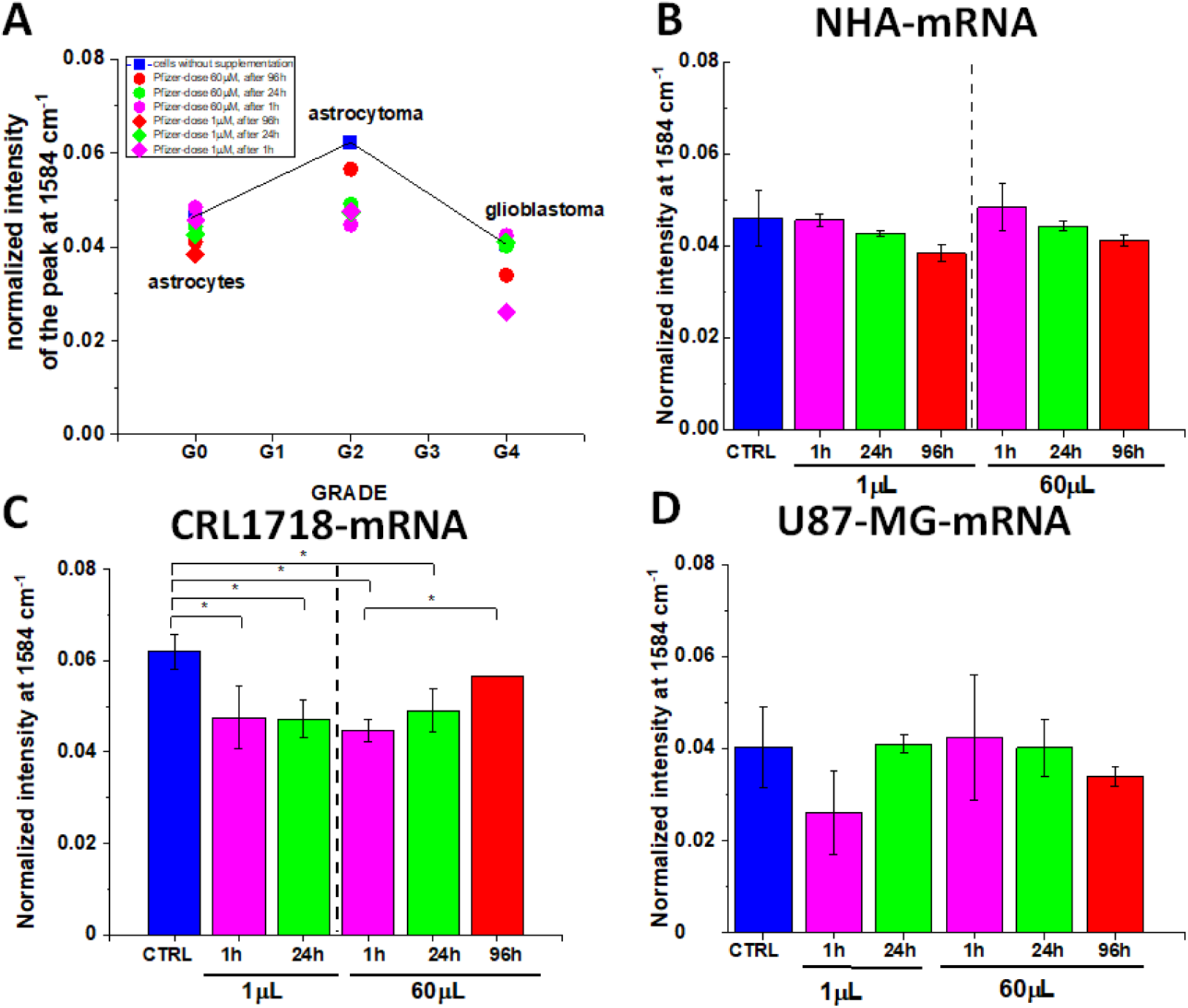
The normalized Raman intensity of the band 1584 cm^-1^, (based on the Raman spectra normalized by vector norm) obtained for mitochondria in normal cells of astrocyte (NHA) (A,B), astrocytoma (CRL-1718) (A,C) and glioblastoma cells (U87 MG)(A,D) without Pfizer/BioNT vaccine (control, blue) and with Pfizer/BioNT vaccine: doses 1 µL/mL and 60 µL/mL, time of incubation 1h-magenta, time of incubation 24h-green, time of incubation 96h-red. The one-way ANOVA using the Tukey test was used to calculate the value significance, asterisk * denotes that the differences are statistically significant, p-value ≤ 0.05.

One can see that the normalized Raman intensity of the band at 1584 cm^-1^ corresponding to the concentration of cytochrome c in mitochondria of cells in vitro decreases upon incubation with mRNA vaccination when compared with the control samples. This effect depends on brain tumor aggressiveness. The observations illustrated in Figure 5 confirm that incubation with the mRNA vaccine decreases the cytochrome c concentration for astrocytoma CRL1718 with statistical significance. Lower concentration of cytochrome c upon incubation with mRNA observed for astrocytoma leads to reduction in mitochondrial membrane potential, reduction of oxidative phosphorylation (respiration) and apoptosis and lessened ATP production.[16,17]

The results presented in Fig. 5 suggest that the Pfizer/BioNT vaccine against COVID-19 reprograms innate immune responses by downregulation of cytochrome c. This conclusion is based on the observation of the cytochrome c may play the role of a universal DAMP molecules (Damage-associated molecular patterns) able of alarming the immune system for danger in any type of cell or tissue and contributing to the host’s defense [21]. Damage-associated molecular patterns (DAMPs) are endogenous danger molecules that are released from damaged or dying cells and activate the innate immune system by interacting with pattern recognition receptors (PRRs).

Our results presented so far fully support these suggestions [17,21]. Cytochrome c is a key protein that is needed to maintain life (respiration via oxidative phosphorylation) and cell death (apoptosis). As a result cytochrome c is a key protein in cancer development. The pandemics has witnessed an explosion in research examining the interplay between the immune response and the intracellular metabolic pathways that mediate it. Research in the field of immunometabolism has revealed that similar mechanisms regulate the host response to infection, autoimmunity, and cancer. Our results suggest that the triangle: cytochrome c- cancer - the immune system are strongly linked.

We found that there is very close relations between cytochrome c and immune system through retinoic acid [22]. Retinoic acid (RA) is an essential molecule in the innate immune system that does not stimulate its ATPase and leads to lack of cytokine induction. Fig.6 shows effect of retinoic acid on cytochrome c in U87 MG cells of glioblastoma.

**Figure 6.**
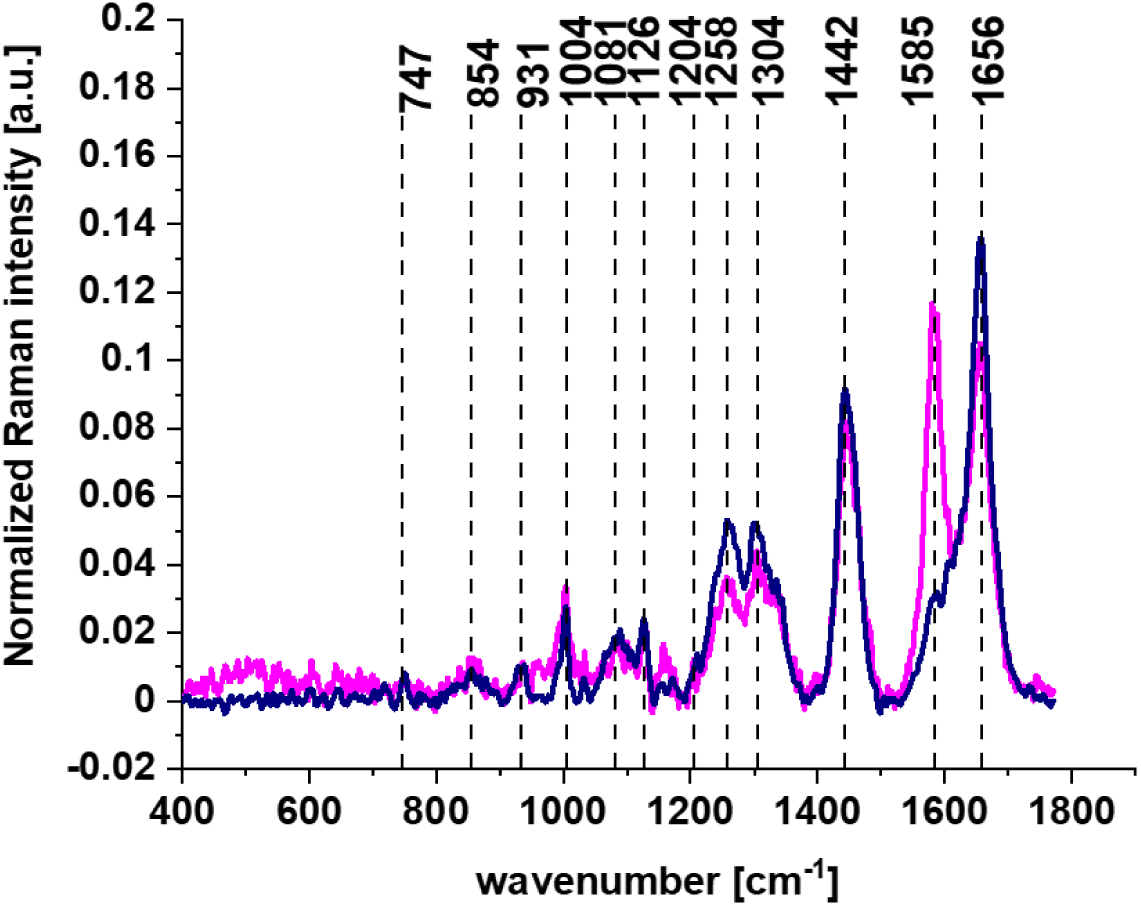
Average Raman spectra for mitochondria for glioblastoma U87-MG cells without (violet) and upon incubation for 24h with retinoic acid (c= 50 µM, pink), (alle spectra represent the arithmetic mean of 3 spectra characteristic for mitochondria of analyzed cells, calculated based on the minimum 1600 single Raman spectra, excitation wavelength 532 nm, laser power 10 mW, integration time 1.0 sec).

Detailed inspection into Figure. 6 demonstrates that the most significant changes occurs at 1584 cm^-1^ corresponding to reduced form of cytochrome c. The Raman intensity of cytochrome c drastically increases upon incubation of U87 MG cells with retinoic acid and represents the reduced form [16,17] in contrast to the oxidized form observed without RA incubation. Incubation in vitro with retinoic acid increases the amount of reduced form of cytochrome c.

Our results from Fig. 6 support the conclusion that retinoic acid is a key player in immunity [23]. Moreover, it has been shown recently that RIG-I (retinoic acid induced gen) triggers a signaling-abortive anti-COVID-19 defense in human lung cells [24].

### 3.3. Nucleus -mRNA

Most scientists claim that mRNA vaccine never enters the nucleus of the cell, which is where our DNA (genetic material) is kept [3]. The cell breaks down and gets rid of the mRNA soon after it is finished using the instructions. It is assumed that it is unplausible that an RNA vaccine would change our DNA, for many reasons. First, neither the coronavirus nor the RNA vaccines (which only code for the spike protein) have a reverse transcriptase. Therefore, RNA vaccines cannot produce DNA molecules. Another reason is that the cells keep their compartments well separated, and messenger RNAs cannot travel from the cytoplasm to the nucleus. Therefore, the vaccine’s mRNA cannot get even close to the DNA, let alone change it. Also, mRNAs are short-lived molecules. The vaccine’s messenger does not stay inside the cell indefinitely, but is degraded after a few hours, without leaving a trace. Finally, clinical studies on thousands of people who have received the RNA vaccines have shown no sign of DNA modification so far. This is perhaps the best indication that these vaccines do not alter our genome.

Therefore is particularly important to monitor alterations in nucleus upon incubation with mRNA by Raman imaging. Figure 7 shows results obtained for nucleus without and upon incubation with vaccine by Raman spectroscopy and imaging.

**Figure 7.**
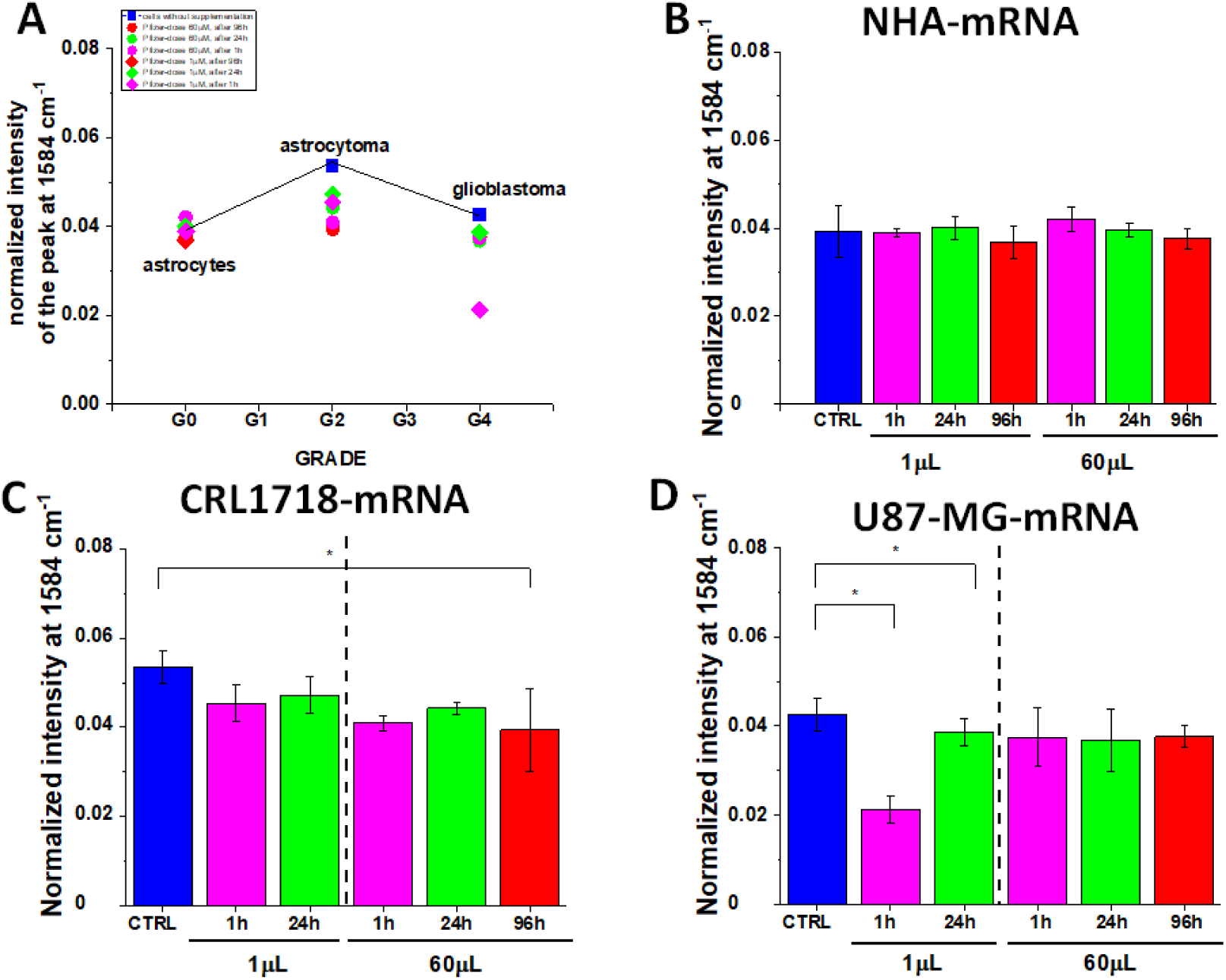
The normalized Raman intensity of the band 1584 cm^-1^, (based on the Raman spectra normalized by vector norm) obtained for nucleus in normal cells of astrocyte (NHA) (A,B), astrocytoma (CRL-1718) (A,C) and glioblastoma cells (U87 MG)(A,D) without Pfizer/BioNT vaccine (control, blue) and with Pfizer/BioNT vaccine: doses 1 µL/mL and 60 µL/mL, time of incubation 1h-magenta, time of incubation 24h-green, time of incubation 96h-red. The one-way ANOVA using the Tukey test was used to calculate the value significance, asterisk * denotes that the differences are statistically significant, p-value ≤ 0.05.

Generally, our results seems to support the conclusions that mRNA vaccine does not enter the DNA of the cell. The Figure. 7 shows that there is no statistical significance for cytochrome c activity for normal astrocytes NHA and U87 MG at 60 µL/mL dose. In contrast, one can observe some statistically significant changes for astrocytoma for the dose 60 µL/mL for the incubation time 96h, and for U87-MG glioblastoma cells for low dose 1 µL/mL for 24h. Because it is believed that mRNA vaccine does not introduce any changes corresponding to DNA, we interpret this result as posttranslational changes in histones, not in DNA. It may indicate that posttranslational changes in histones of the nucleus occur upon incubation with the mRNA vaccine.

### 3.4. Lipid droplets-mRNA and rough endoplasmic reticulum

The significant changes in cytochrome c concentration occurs not only for mitochondria. Similar alterations in biochemical composition of cytochrome c reflected by the band at 1584 cm^-1^ are also observed in lipid droplets and in lipid structures of rough endoplasmic reticulum (RER) (Figures 8 and 9). RER is studded with ribosomes that perform biological protein synthesis (mRNA translation). Ribosomes are the sites of protein synthesis for mRNA vaccines. Ribosomes are too small to be seen by resolution of Raman imaging.

**Figure 8.**
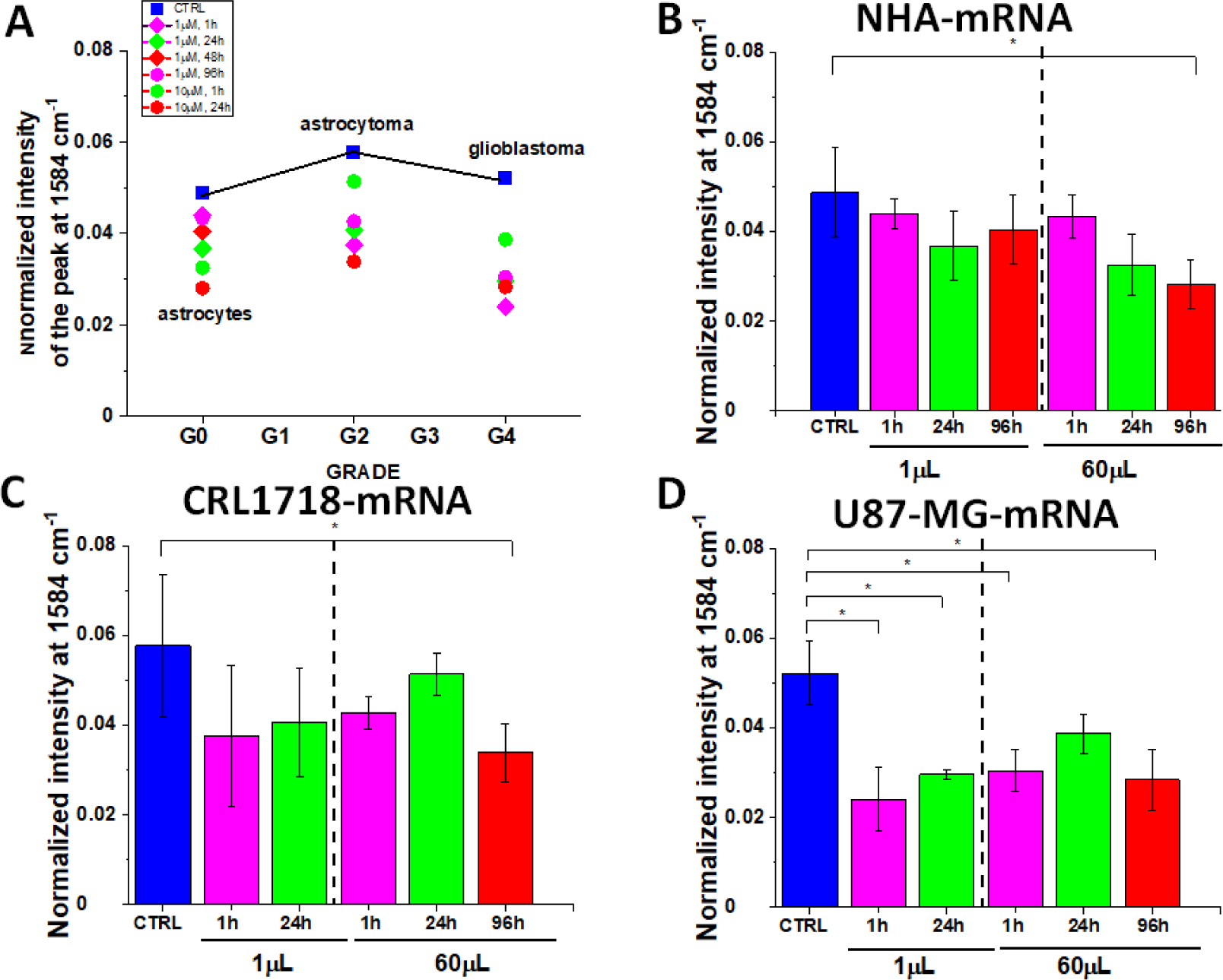
The normalized Raman intensity of the band 1584 cm^-1^, (based on the Raman spectra normalized by vector norm) obtained for lipid droplets in normal cells of astrocyte (NHA) (A,B), astrocytoma (CRL-1718) (A,C) and glioblastoma cells (U87 MG)(A,D) without Pfizer/BioNT vaccine (control, blue) and with Pfizer/BioNT vaccine: doses 1 µL/mL and 60 µL/mL, time of incubation 1h-magenta, time of incubation 24h-green, time of incubation 96h-red. The one-way ANOVA using the Tukey test was used to calculate the value significance, asterisk * denotes that the differences are statistically significant, p-value ≤ 0.05.

**Figure 9.**
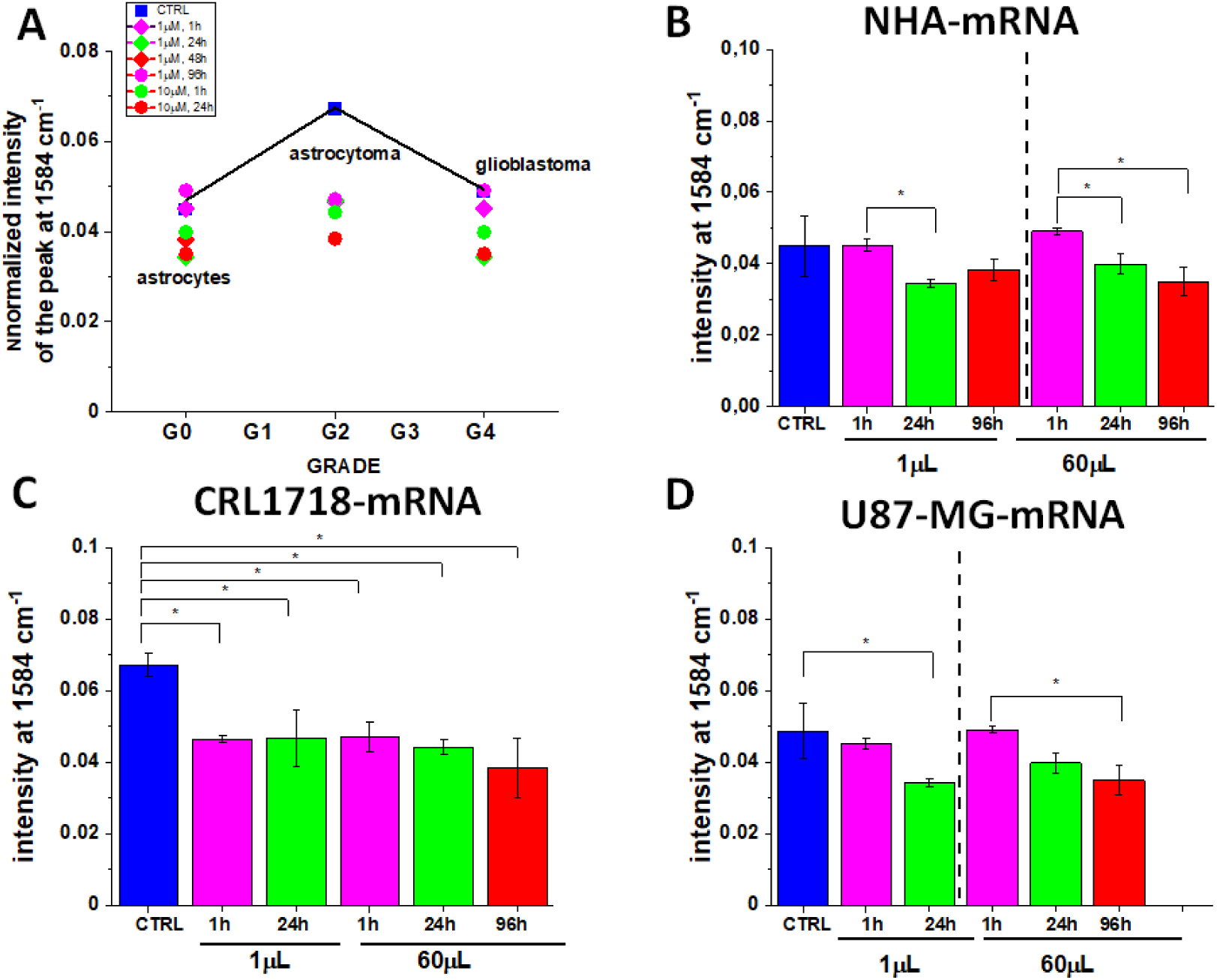
The normalized Raman intensity of the band 1584 cm^-1^, (based on the Raman spectra normalized by vector norm) obtained for rough endoplasmatic reticulum in normal cells of astrocyte (NHA) (A,B), astrocytoma (CRL-1718) (A,C) and glioblastoma cells (U87 MG)(A,D) without Pfizer/BioNT vaccine (control, blue) and with Pfizer/BioNT vaccine: doses 1 µL/mL and 60 µL/mL, time of incubation 1h-magenta, time of incubation 24h-green, time of incubation 96h-red. The one-way ANOVA using the Tukey test was used to calculate the value significance, asterisk * denotes that the differences are statistically significant, p-value ≤ 0.05.

The Figure. 8 and 9 shows that the cytochrome c signal in lipid droplets and rough endoplasmatic reticulum at 1584 cm^-1^ decreases for all types of studied glial cells: normal cells, astrocytoma, and glioblastoma, all periods of incubation and doses with statistical significance.

This trends in lipid droplets and rough endoplasmatic reticulum correlate with changes observed in mitochondria (Fig.5).

The alterations in lipid composition can also be monitored by the band at 2845 cm^-1^ that monitor concentration of triglycerides (TAG) [22,25,26]. Figure. 9 shows a comparison of the average Raman spectra (normalized by vector norm) obtained from the cluster analysis for mitochondria at 532 nm excitation with and without mRNA vaccine in the high frequency spectral region.

Recently we found that the Raman signal intensity of the band at 2845 cm^-1^ in lipid droplets is significantly higher for high-grade cancer of glioblastoma (U87 MG) than for normal astrocytes (NHA), indicating a higher concentration of TAGs in cancer lipid droplets [20]. The higher concentration of TAG was related to the increased amount of cytoplasmic lipid droplets in human glioblastoma cells in comparison to normal astrocytes.

Recently we showed that the lipid droplets in cancer cells are predominantly filled with TAGs and are involved in energy storage. The lipid droplets in normal cells NHA are filled mainly with retinyl esters /retinol and are involved in signaling, especially JAK2/STAT6 pathway signaling [22]. The results presented in Figure 9 suggest that upon incubation with mRNA increases the role of signaling. Our results support those reported recently that COVID-19 spike protein elicits cell signaling in human host cells, which may have serious implications for Possible Consequences of COVID-19 vaccines [27].

### 3.5. Cytoplasm-mRNA

Vaccine mRNA is translated by ribosomes in the cytoplasm. Therefore is particularly important to monitor alterations in cytoplasm upon incubation with mRNA. Figure. 11 shows effect of incubation with mRNA compared to the control cells without mRNA in cytoplasm.

One can see that cytochrome c activity at 1584 cm^-1^ in cytoplasm increases upon incubation with mRNA for normal astrocytes (NHA) at 60 µL/mL and for glioblastoma U87 MG at 1µL/mL. In contrast, for astrocytoma and glioblastoma U87 MG The Raman signal at 1584 cm^-1^ decreases upon incubation with mRNA. Statistical significance is presented in Figures 10 B,C,D.

**Figure 10.**
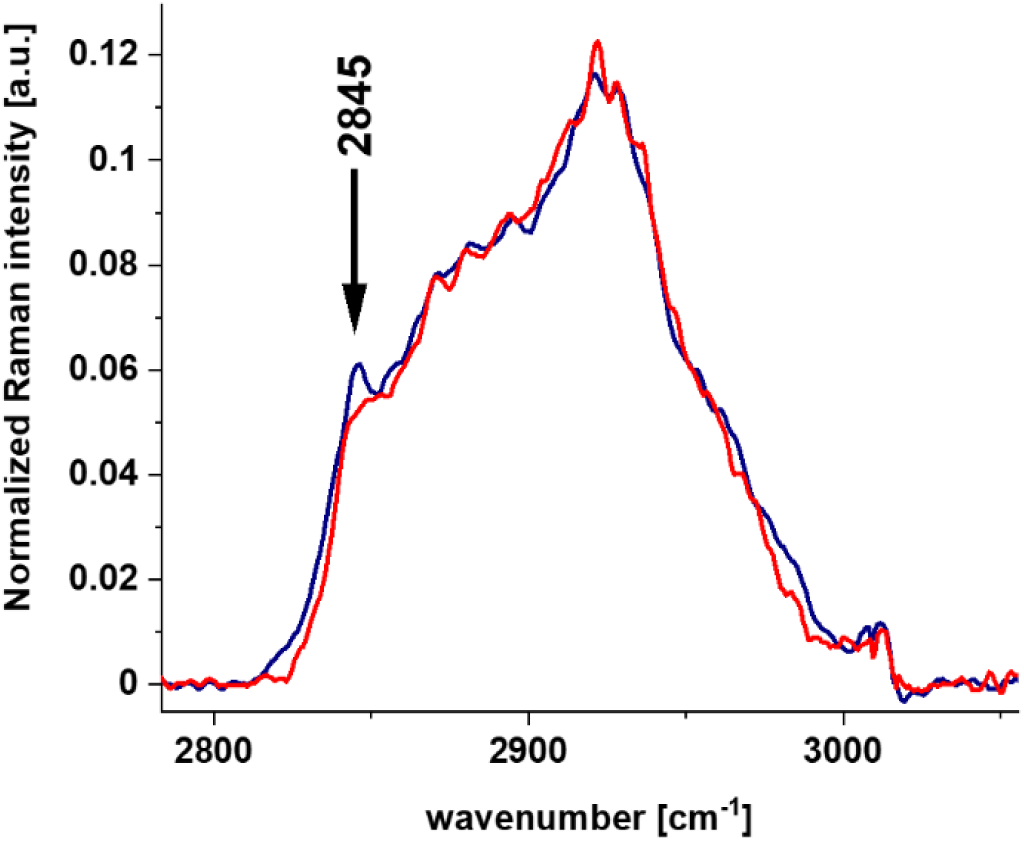
The effect of the Pfizer/BioNT vaccine on lipid droplets in glioblastoma cells (U87 MG) by Raman imaging: for U87-MG control cells without vaccine (dark blue) and upon incubation with vaccine for 96h for the dose 60 µL/mL (red).

The release of Cyt c from mitochondria to the cytosol activates apoptosis by the stream of caspase and proteases.[28] The concept that programmed cell death by apoptosis serves as a natural barrier to cancer development.[29] Therefore, the results provide new tools to check the role of Cyt c in an anti-apoptotic switch by phosphorylation of Tyr48.[30]

Our results from Fig. 10 show that apoptosis is reduced upon mRNA vaccine for astrocytoma and glioblastoma which indicate decreased ability to fight against cancer development by programmed cell death.

### 3.6. Membrane-mRNA

As it is well known professional antigen-presenting cells (APCs) play a crucial role in initiating immune responses. Under pathological conditions also epithelial cells act as nonprofessional APCs, thereby regulating immune responses at the site of exposure. Therefore it is interesting to monitor alterations at the surface of the cell membranes upon incubation with mRNA.

Figure. 12 shows effect of incubation with mRNA compared to the control cells without mRNA at surface of the membrane.

**Figure 11.**
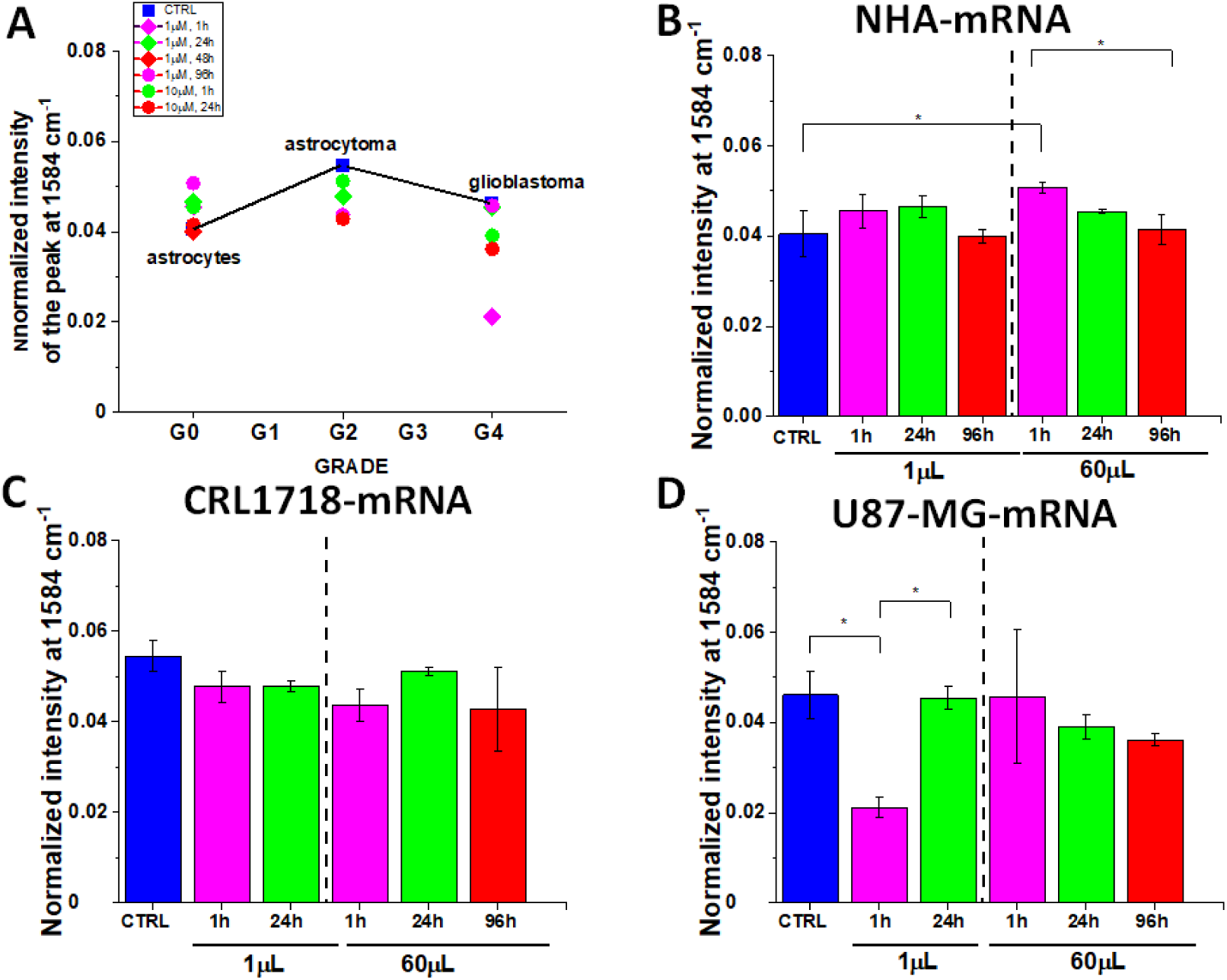
The normalized Raman intensity of the band 1584 cm^-1^, (based on the Raman spectra normalized by vector norm) obtained for cytoplasm in normal cells of astrocyte (NHA) (A,B), astrocytoma (CRL-1718) (A,C) and glioblastoma cells (U87 MG)(A,D) without Pfizer/BioNT vaccine (control, blue) and with Pfizer/BioNT vaccine: doses 1 µL/mL and 60 µL/mL, time of incubation 1h-magenta, time of incubation 24h-green, time of incubation 96h-red. The one-way ANOVA using the Tukey test was used to calculate the value significance, asterisk * denotes that the differences are statistically significant, p-value ≤ 0.05.

**Figure 12.**
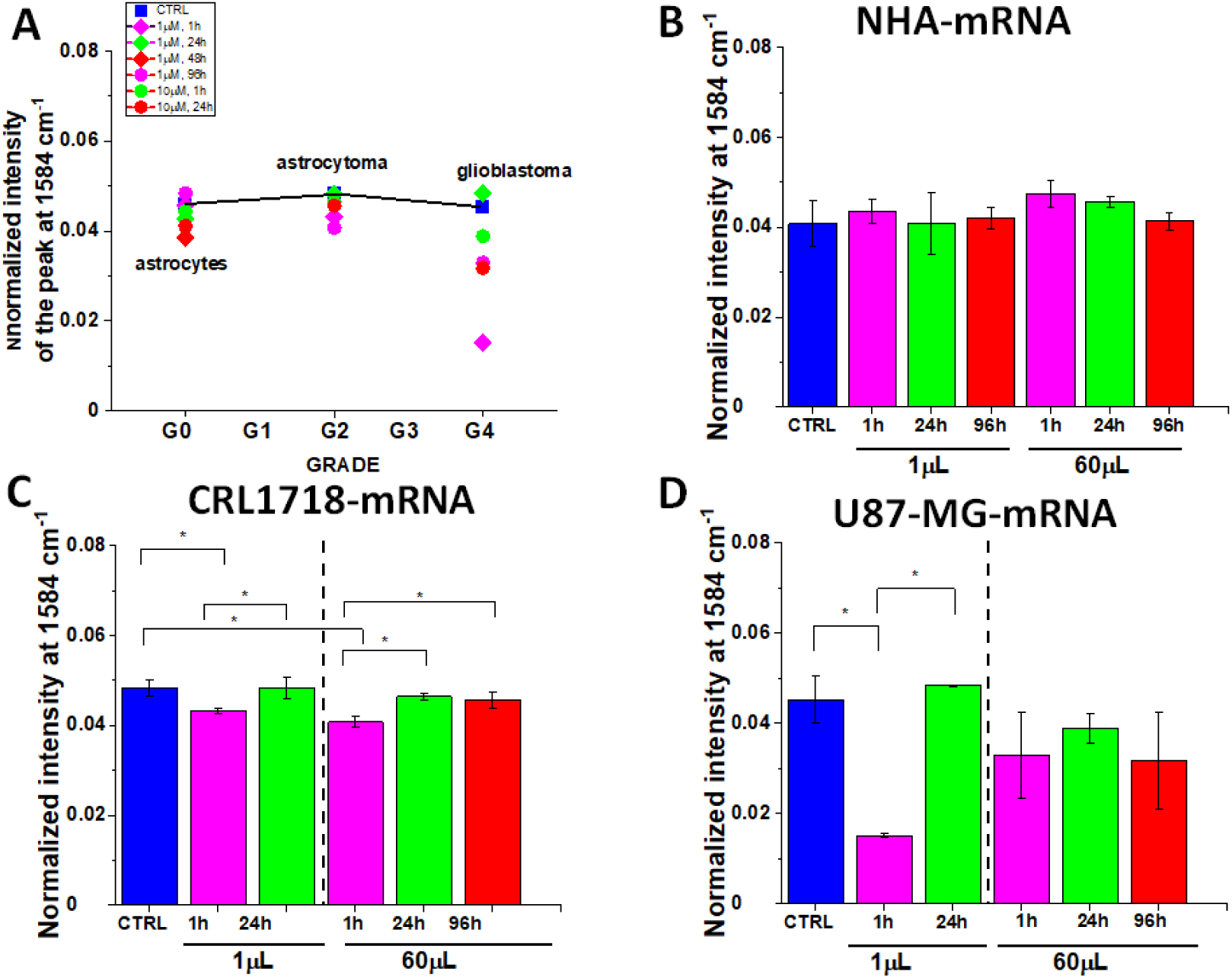
The normalized Raman intensity of the band 1584 cm^-1^, (based on the Raman spectra normalized by vector norm) obtained for cell membrane in normal cells of astrocyte (NHA) (A,B), astrocytoma (CRL-1718) (A,C) and glioblastoma cells (U87 MG)(A,D) without Pfizer/BioNT vaccine (control, blue) and with Pfizer/BioNT vaccine: doses 1 µL/mL and 60 µL/mL, time of incubation 1h-magenta, time of incubation 24h-green, time of incubation 96h-red. The one-way ANOVA using the Tukey test was used to calculate the value significance, asterisk * denotes that the differences are statistically significant, p-value ≤ 0.05.

One can see that cytochrome c activity at 1584 cm^-1^ decreases for glioblastoma U87 MG with statistical significance at p-value ≤ 0.05.

Table 1. summarize the results presented above for the Raman intensity band at 1584 cm^-1^ for mitochondria, nucleus lipid droplets, cytoplasm and cell membrane for NHA, CRL1718 and U-87 MG cell lines, presented as the mean ± SD based on the normalized Raman spectra. (normalization: divided by norm).

**Table 1.** The comparison of the Raman intensity band at 1584 cm^-1^ for mitochondria, nucleus, lipid droplets, cytoplasm and cell membrane for NHA, CRL1718 and U-87 MG cell lines, presented as the mean ± SD based on the normalized Raman spectra (normalization: divided by norm). Raman bands intensity were taken from normalized by vector norm spectra, number of cells=3.

|  | Dose | Incubation<br>time | MITOCHONDRIA | NUCLEUS | LIPID<br>DROPLETS | CYTOPLASM | MEMBRANE |
| --- | --- | --- | --- | --- | --- | --- | --- |
| NHA |  |  | 0.04607±0.00613 | 0.0393±0.00591 | 0.04865±0.01003 | 0.04053±0.00515 | 0.04076±0.00515 |
|  | mRNA<br>60 µL/mL | 1h | 0.04836±0.00512 | 0.04208±0.00287 | 0.04337±0.00476 | 0.05073±0.00107 | 0.04742±0.00287 |
|  |  | 24h | 0.04435±0.00104 | 0.03966±0.00154 | 0.03254±0.00693 | 0.04545±4.60241E-4 | 0.04566±0.0012 |
|  |  | 96h | 0.04112±0.00127 | 0.03762±0.00233 | 0.02808±0.0054 | 0.04154±0.00332 | 0.04136±0.002 |
|  | 1 µL/mL | 1h | 0.04567±0.00132 | 0.0389±0.001 | 0.04387±0.00344 | 0.0455±0.00387 | 0.04355±0.00267 |
|  |  | 24h | 0.04262±5.7E-4 | 0.04005±0.00267 | 0.03668±0.00773 | 0.04657±0.00235 | 0.0407±0.0069 |
|  |  | 96h | 0.0384±0.00182 | 0.03693±0.00371 | 0.04037±0.00766 | 0.03996±0.0016 | 0.04206±0.00246 |
| CRL1718 |  |  | 0.06203±0.00381 | 0.05355±0.00364 | 0.05772±0.01587 | 0.05456±0.00338 | 0.04844±0.0018 |
|  | mRNA<br>60 µL/mL | 1h | 0.04469±0.00243 | 0.04091±0.00178 | 0.04265±0.00358 | 0.04368±0.00364 | 0.04071±0.00121 |
|  |  | 24h | 0.04908±0.00479 | 0.04429±0.00149 | 0.05135±0.00475 | 0.05119±9.37692E-4 | 0.04637±6.53265E-4 |
|  |  | 96h | 0.05654±6E-5 | 0.03941±0.0093 | 0.03382±0.00646 | 0.04282±0.00928 | 0.04566±0.00176 |
|  | 1 µL/mL | 1h | 0.04757±0.00683 | 0.04544±0.00416 | 0.03745±0.01578 | 0.04775±0.00352 | 0.04323±5.60139E-4 |
|  |  | 24h | 0.04723±0.00405 | 0.04723±0.00405 | 0.04069±0.01214 | 0.04778±0.00121 | 0.0483±0.0024 |
| U87-MG |  |  | 0.03106±0.0075 | 0.03054±0.00809 | 0.01434±0.00424 | 0.0326±0.00659 | 0.04011±0.00312 |
|  | mRNA<br>60 µL/mL | 1h | 0.04242±0.01364 | 0.03744±0.00657 | 0.03042±0.00471 | 0.04573±0.01483 | 0.03293±0.0096 |
|  |  | 24h | 0.04014±0.00625 | 0.03682±0.00696 | 0.03869±0.00443 | 0.03911±0.00275 | 0.03883±0.00339 |
|  |  | 96h | 0.03397±0.0022 | 0.03761±0.00247 | 0.02832±0.00686 | 0.03616±0.00138 | 0.03174±0.01082 |
|  | 1 µL/mL | 1h | 0.02613±0.00916 | 0.02123±0.00296 | 0.02406±0.00701 | 0.02121±0.00215 | 0.0151±5.06976E-4 |
|  |  | 24h | 0.04102±0.00192 | 0.03867±0.0031 | 0.02953±9.76761E-4 | 0.04547±0.00243 | 0.04836±1.64306E-4 |

## 4. Conclusions

We showed that new tools of Raman imaging we present in this paper raise exciting possibilities for new ways to understand links between pathways of cancer, immune responses, and recognize metabolites that regulates these pathways.

We used Raman spectroscopy to monitor changes in the redox state of the mitochondrial cytochromes in human brain cells *in vitro* of normal astrocytes, astrocytoma, glioblastoma upon incubation with mRNA vaccine. We observed the effect of the mRNA vaccine on biodistribution of different chemical components, particularly cytochrome c, in the specific organelles of human brain glial cells: nucleus, mitochondria, lipid droplets, cytoplasm, rough endoplasmatic reticulum and membrane.

We showed that mRNA vaccine (Pfizer) changes mitochondria by downregulation of cytochrome c resulting in lower effectiveness of respiration (oxidative phosphorylation) and lower ATP production. It can lead to lower immune system response.

Decrease of Amide I concentration in mitochondrial membrane potential may suggest functional deterioration of the adenine nucleotide translocator. mRNA vaccine modifies significantly de novo lipids synthesis in lipid droplets. The results presented in paper suggest that upon incubation with mRNA the role of signaling function of lipid droplets increases. The observed alterations in biochemical profiles upon incubation with the Pfizer/BioNT in the specific organelles of the glial cells are similar to those we observe for brain cancer vs grade of aggressiveness.

## Author Contributions

Conceptualization-H.A.; funding acquirement- H.A.; investigation- B.B-P, K.B.; visualization- B. B-P.; writing-original draft - H.A., B. B-P.; writing - B.B-P., K.B.; review - H.A; editing-B.B-P., K.B. All authors have read and agreed to the published version of the manuscript.

## Data Availability Statement

The raw data underlying the results presented in the study are available from Lodz University of Technology Institutional Data Access. Request for access to those data should be addressed to the Head of Laboratory of Laser Molecular Spectroscopy, Institute of Applied Radiation Chemistry, Lodz University of Technology. Data requests might be sent by email to the secretary of the Institute of Applied Radiation Chemistry.

## References

1. WHO Coronavirus (COVID-19) Dashboard | WHO Coronavirus (COVID-19) Dashboard With Vaccination Data Available online: https://covid19.who.int/ (accessed on 12 October 2021).

2. Doshi, P. Covid-19 Vaccines: In the Rush for Regulatory Approval, Do We Need More Data? BMJ 2021, 373, 2020–2022.

3. Bettini, E.; Locci, M. SARS-CoV-2 MRNA Vaccines: Immunological Mechanism and Beyond. Vaccines 2021, 9, 147.

4. Ols, S.; Yang, L.; Thompson, E.A.; Pushparaj, P.; Tran, K.; Liang, F.; Lin, A.; Eriksson, B.; Karlsson Hedestam, G.B.; Wyatt, R.T.; et al. Route of Vaccine Administration Alters Antigen Trafficking but Not Innate or Adaptive Immunity. Cell Rep. 2020, 30, 3964-3971.e7.

5. Bahl, K.; Senn, J.J.; Yuzhakov, O.; Bulychev, A.; Brito, L.A.; Hassett, K.J.; Laska, M.E.; Smith, M.; Almarsson, Ö.; Thompson, J.; et al. Preclinical and Clinical Demonstration of Immunogenicity by MRNA Vaccines against H10N8 and H7N9 Influenza Viruses. Mol. Ther. 2017, 25, 1327.

6. Walt, D.R.; Ogata, A.F.; Cheng, C.-A.; Desjardins, M.; Senussi, Y.; Sherman, A.C.; Powell, M.; Novack, L.; Von, S.; Li, X.; et al. Circulating Severe Acute Respiratory Syndrome Coronavirus 2 (SARS-CoV-2) Vaccine Antigen Detected in the Plasma of MRNA-1273 Vaccine Recipients. Clin. Infect. Dis. 2021.

7. Rhea, E.M.; Logsdon, A.F.; Hansen, K.M.; Williams, L.M.; Reed, M.J.; Baumann, K.K.; Holden, S.J.; Raber, J.; Banks, W.A.; Erickson, M.A. The S1 Protein of SARS-CoV-2 Crosses the Blood–Brain Barrier in Mice. Nat. Neurosci. 2020 243 2020, 24, 368–378.

8. YC, L.; WZ, B.; T, H. The Neuroinvasive Potential of SARS-CoV2 May Play a Role in the Respiratory Failure of COVID-19 Patients. J. Med. Virol. 2020, 92, 552–555.

9. K, S.; M, B.; A, S.; N, R. The Involvement of the Central Nervous System in Patients with COVID-19. Rev. Neurosci. 2020, 31, 453–456.

10. Haider, A.; Siddiqa, A.; Ali, N.; Dhallu, M. COVID-19 and the Brain: Acute Encephalitis as a Clinical Manifestation. Cureus 2020, 12.

11. K, S.; J, M.; L, V.; G, F.; S, F.; K, T.; M, P.; WJ, S.; S, Z.; B, M. Prior Infection and Passive Transfer of Neutralizing Antibody Prevent Replication of Severe Acute Respiratory Syndrome Coronavirus in the Respiratory Tract of Mice. J. Virol. 2004, 78, 3572–3577.

12. J, X.; S, Z.; J, L.; L, L.; Y, L.; X, W.; Z, L.; P, D.; J, Z.; N, Z.; et al. Detection of Severe Acute Respiratory Syndrome Coronavirus in the Brain: Potential Role of the Chemokine Mig in Pathogenesis. Clin. Infect. Dis. 2005, 41, 1089–1096.

13. CK, B.; HT, H.; A, B.-T.; WR, B.; T, H.; C, K.; MM, L.; A, P.; JC, R.; FCM, S.; et al. Infectability of Human BrainSphere Neurons Suggests Neurotropism of SARS-CoV-2. ALTEX 2020, 37, 665–671.

14. Crunfli, F.; Carregari, V.C.; Veras, F.P.; Vendramini, P.H.; Valença, A.G.F.; Antunes, A.S.L.M.; Brandão-Teles, C.; Zuccoli, G. da S.; Reis-de-Oliveira, G.; Silva-Costa, L.C.; et al. SARS-CoV-2 Infects Brain Astrocytes of COVID-19 Patients and Impairs Neuronal Viability. medRxiv 2020, 18, 2020.10.09.20207464.

15. Ndeupen, S.; Qin, Z.; Jacobsen, S.; Estanbouli, H.; Bouteau, A.; Igyártó, B.Z. The MRNA-LNP Platform’s Lipid Nanoparticle Component Used in Preclinical Vaccine Studies Is Highly Inflammatory. bioRxiv 2021, 2021.03.04.430128.

16. Abramczyk, H.; Surmacki, J.M.; Brozek-Pluska, B.; Kopec, M. Revision of Commonly Accepted Warburg Mechanism of Cancer Development: Redox-Sensitive Mitochondrial Cytochromes in Breast and Brain Cancers by Raman Imaging. Cancers 2021, Vol. 13, Page 2599 2021, 13, 2599.

17. Abramczyk, H.; Brozek-Pluska, B.; Kopec, M.; Surmacki, J.; Błaszczyk, M.; Radek, M. Redox Imbalance and Biochemical Changes in Cancer by Probing Redox-Sensitive Mitochondrial Cytochromes in Label-Free Visible Resonance Raman Imaging. Cancers (Basel). 2021, 13, 960.

18. Comirnaty (COVID-19 Vaccine, MRNA) 4 Dosage And Administration | Pfizer Medical Information - Canada Available online: https://www.pfizermedicalinformation.ca/en-ca/pfizer-biontech-covid-19-vaccine/4-dosage-and-administration (accessed on 2 March 2022).

19. Vaughn, A.E.; Deshmukh, M. Glucose Metabolism Inhibits Apoptosis in Neurons and Cancer Cells by Redox Inactivation of Cytochrome C. Nat. Cell Biol. 2008, 10, 1477.

20. Atlante, A.; Valenti, D. A Walk in the Memory, from the First Functional Approach up to Its Regulatory Role of Mitochondrial Bioenergetic Flow in Health and Disease: Focus on the Adenine Nucleotide Translocator. Int. J. Mol. Sci. 2021, Vol. 22, Page 4164 2021, 22, 4164.

21. Eleftheriadis, T.; Pissas, G.; Liakopoulos, V.; Stefanidis, I. Cytochrome c as a Potentially Clinical Useful Marker of Mitochondrial and Cellular Damage. Front. Immunol. 2016, 7, 279.

22. Abramczyk, H.; Imiela, A.; Surmacki, J. Novel Strategies of Raman Imaging for Monitoring Intracellular Retinoid Metabolism in Cancer Cells. J. Mol. Liq. 2021, 334, 116033.

23. Pino-Lagos, K.; Guo, Y.; Noelle, R.J. Retinoic Acid: A Key Player in Immunity. BioFactors 2010, 36, 430–436.

24. Yamada, T.; Sato, S.; Sotoyama, Y.; Orba, Y.; Sawa, H.; Yamauchi, H.; Sasaki, M.; Takaoka, A. RIG-I Triggers a Signaling-Abortive Anti-SARS-CoV-2 Defense in Human Lung Cells. Nat. Immunol. 2021 227 2021, 22, 820–828.

25. Abramczyk, H.; Brozek-Pluska, B.; Jarota, A.; Surmacki, J.; Imiela, A.; Kopec, M. A Look into the Use of Raman Spectroscopy for Brain and Breast Cancer Diagnostics: Linear and Non-Linear Optics in Cancer Research as a Gateway to Tumor Cell Identity. Expert Rev. Mol. Diagn. 2020, 20, 99–115.

26. DL, B.; G, D.; L, S.; R, W. Proteomic Analysis of Proteins Associated with Lipid Droplets of Basal and Lipolytically Stimulated 3T3-L1 Adipocytes. J. Biol. Chem. 2004, 279, 46835–46842.

27. Suzuki, Y.J.; Gychka, S.G. SARS-CoV-2 Spike Protein Elicits Cell Signaling in Human Host Cells: Implications for Possible Consequences of COVID-19 Vaccines. Vaccines 2021, 9, 1–8.

28. Kroemer, G.; Dallaporta, B.; Resche-Rigon, M. The Mitochondrial Death/Life Regulator in Apoptosis and Necrosis. Annu. Rev. Physiol. 1998, 60, 619–642.

29. Hanahan, D.; Weinberg, R.A. Hallmarks of Cancer: The next Generation. Cell 2011, 144, 646–674.

30. Hüttemann, M.; Pecina, P.; Rainbolt, M.; Sanderson, T.H.; Kagan, V.E.; Samavati, L.; Doan, J.W.; Lee, I. The Multiple Functions of Cytochrome c and Their Regulation in Life and Death Decisions of the Mammalian Cell: From Respiration to Apoptosis. Mitochondrion 2011, 11, 369–381.

